# Global structure of new relational knowledge networks is represented in retrosplenial complex, and node-distance in hippocampus

**DOI:** 10.64898/2026.07.24.740285

**Authors:** Gordon B. Feld, Pablo F. Velasco, Christoffer Gahnstrom, Martin F. Gerchen, Qasim Mian, Matthieu Bernard, Mona M. Garvert, Fred Dick, Hugo J. Spiers

## Abstract

Much of our knowledge is structured in networks, but how the brain represents such networks remains unclear. In most nested knowledge structures a subset of the nodes will be highly connected to other nodes in the network. High levels of connection can exist either locally (high degree centrality) or at a global level (high closeness/betweenness centrality). Here, we explored how the human brain represents network structure using functional magnetic resonance imaging (fMRI) and a novel learning paradigm. During training, participants learned the transition structure for travel between a set of fictious partially connected alien planets. Despite encountering only individual connections, participants choices indicated they could infer the broader network structure. Next day, they viewed each stimulus during a cover task while undergoing fMRI. Representational-similarity analysis showed that shortest-path distances within the learned graph were encoded in the posterior hippocampus and right retrosplenial complex, whereas global connectivity (closeness/ betweenness centrality) was represented in the left retrosplenial complex. These findings extend the view that brain networks associated with spatial navigation also process abstract relational knowledge using principles akin to mapping physical space. The results are also consistent with proposals that the hippocampus represents distance information and that the retrosplenial cortex acts as hub for integrating recently acquired knowledge.

## Introduction

The German philosopher Immanuel Kant argued that humans employ their capacity for navigation not only in physical space but also in the space of thought: when we think, we are navigating conceptual spaces (Kant, 1786). The intimate connection between spatial and abstract knowledge spaces is evident in our abundant use of spatial terms to describe abstract entities (Lakoff & Johnson, 2008; Landau & Jackendoff, 1993), as well as in mnemonic techniques the world over — from the ancient Roman method of building a mental memory palace to the songlines of aboriginal Australia or the memory tablets of the Luba people (Fernandez-Velasco & Spiers, 2024; Norris & Harney, 2014; Roberts & Roberts, 1996; Yates, 2013). More recently, this connection has also garnered support from evidence that the same neural substrates that store cognitive maps of physical environments also represent abstract knowledge spaces (Bellmund et al., 2018; Epstein et al., 2017).

Many animals appear to construct an allocentric representation of space – a cognitive map – from a sequence of egocentric experiences. This map-like representation is thought to strongly depend on neurons in the hippocampal and parahippocampal regions which have been found in rodents to correlate with measures of place, distance travelled, head-direction and proximity to boundaries (Hafting et al., 2005; Hoydal et al., 2019; Lever et al., 2009; O’Keefe & Dostrovsky, 1971; O’Keefe & Nadel, 1978; Solstad et al., 2008; Spiers et al., 2018; Taube et al., 1990). In humans such regions have been found to represent distance (Bush et al., 2017; Howard et al., 2014; Huang et al., 2024; Morgan et al., 2011; Patai et al., 2019) and direction information (Chadwick et al., 2015; Patai et al., 2019; Stemmler et al., 2015).

Recent evidence has shown that the same brain structures also support cognitive maps of abstract knowledge spaces, coding for temporal (Bellmund et al., 2019; Bellmund et al., 2022; Solomon et al., 2019), social (Park et al., 2021; Tavares et al., 2015) and conceptual relationships (Constantinescu et al., 2016; Garvert et al., 2017; Morton et al., 2021; Theves et al., 2020; Zheng et al., 2024). This line of work suggests that the cognitive maps supported by hippocampal and parahippocampal regions are domain-general and serve to systematically organise information across both spatial and non-spatial dimensions (Behrens et al., 2018; Spiers, 2020). If this view is correct, then the abstract knowledge stored in cognitive maps may be navigated much like we navigate real spaces (Bellmund et al., 2018; Schafer & Schiller, 2018; Vigano et al., 2023; Vigano et al., 2021). Furthermore, the computations that allow for spatial navigation may be used to navigate abstract knowledge spaces, in line with existing theories linking abstract and spatial reasoning (Hills et al., 2008; Solomon et al., 2019).

One way to explore the connection between space and abstract thought is by considering the network structures in which both spatial and abstract knowledge are organised (George et al., 2021; Mok & Love, 2019; Schapiro et al., 2016; Whittington et al., 2020). For instance, a subway network, such as the London underground, can be thought of as a series of nodes (the stations) and edges (the connections between stations) (Balaguer et al., 2016). Road networks can similarly be represented through graphs, and conventional cartographic maps can be converted to graphs (Pung et al., 2022). Considering the network structures underlying relational knowledge allows us to describe, visualise and analyse this knowledge by using graph theory (Baronchelli et al., 2013; Lynn & Bassett, 2020). And these same theoretical tools can be applied to the study of abstract networks such as semantics (Steyvers & Tenenbaum, 2005), music (Liu et al., 2010), social relations (Barabâsi et al., 2002; Girvan & Newman, 2002), or concepts (Masucci et al., 2011).

In a network, the simplest properties are those of nodes and edges/connections. How many direct neighbours a node has is called degree centrality, which it is considered a measure of the local ‘importance’ of a node. The degree of a node can be used to describe not just street networks, but also social networks (e.g. how many friends a person has; (Barabâsi et al., 2002), or scientific influence (e.g. the number of citations of a paper; (Martin et al., 2013). There are also metrics that relate individual nodes to the global topology of a network. Two commonly considered global metrics are closeness centrality and betweenness centrality, which can be calculated for each node in the network. Closeness centrality is a measure of how many connections must one take to reach all other nodes in the network from a given node. Betweenness centrality is a measure of how often a shortest path of the network passes through a specific node. In a city network the central streets will tend to have high closeness centrality as they are topologically closer to all other streets in the network. By contrast bridges in a city centre will typically have high betweenness centrality because many shortest paths will be forced to cross over such bridges. Both closeness and betweenness centrality provide an indicator of the global ‘importance’ of a given node in the network.

By collecting fMRI data while participants navigated London’s (UK) streets in a film simulation it has been possible to examine how brain regions dynamically respond to the connectivity of a street network (Javadi et al., 2017). Upon entering a new street, activity in the posterior right hippocampus tracked changes in centrality measures. Hippocampal activity increased when the new street had more connections (higher degree centrality) and decreased when it had fewer connections. Such a response is consistent with the idea that the hippocampus may serve to simulate future trajectories in the network of possible future states when a new state is entered (Bendor & Spiers, 2016; Olafsdottir et al., 2015; Widloski & Foster, 2022).

Another measure in networks is link distance, which is the number of nodes between a pair of nodes in a network. Past electrophysiological and fMRI studies have found evidence for activity in the hippocampus tracking the distance to the goal in navigation (Balaguer et al., 2016; Howard et al., 2014; Patai et al., 2019; Sherrill et al., 2013; Spiers & Barry, 2015; Spiers & Maguire, 2007a; Spiers et al., 2018) and distance between landmarks in a city (Morgan et al., 2011). While most studies have used the metric distance (e.g., meters), one fMRI study, using a virtual subway network, found hippocampal activity tracked the link distance from the current node to the final destination node, increasing with proximity (Balaguer et al., 2016). However, because distance was correlated with trial end and reward, it is unclear if this hippocampal response was mainly driven by link distance or expectations in reward, or less cognitive demand in the last approach. Notably the anterior hippocampal response occurred in conjunction with other regions in the default mode network.

While the posterior hippocampus appears to represent information about distance and local connectivity in networks, other brain areas may be important in coding the global connectivity of a network. Three brain regions are plausible candidates: the anterior hippocampus, entorhinal cortex and retrosplenial cortex. All are core regions in the spatial navigation network (Epstein et al., 2017). When entering new streets in London, the right anterior hippocampus was found to track the changes in the closeness centrality of the street network (Javadi et al., 2017), consistent with its argued role in broader, large-scale representations of space (Farzanfar et al., 2023; Poppenk et al., 2013). However, in the real-world streets of London, closeness centrality changes were correlated with changes in the length of the street, because highly connected streets tend to be longer. Thus, it remains unclear if the anterior hippocampal response was driven by the view or network properties. While the entorhinal cortex has not been linked to representing global networks, its activity has been used to reconstruct the structure of a network of learnt individual associations between presented objects (Garvert et al., 2017). Such a reconstruction suggests that the entorhinal cortex might contain more than simply degree centrality information of items. Finally, the retrosplenial cortex has been implicated in integrating and consolidating spatial knowledge (Farzanfar et al., 2023; Spiers & Maguire, 2007b; Vann et al., 2009). Because the global connection structure in the network cannot be directly appreciated from one node, it requires accumulating knowledge over many trials. The retrosplenial cortex, rather than hippocampus, has been found to track the distance to goal locations in familiar environments, where extended exposure to the network structure has occurred (Patai et al., 2019) and to represent distances in social networks (Peer et al., 2021). However, it remains unclear whether this extends to networks of newly learned abstract information.

To determine whether different brain regions represent distance and centrality metrics we designed a graph-learning task with a topology that allows us to disentangle contributions of local and global connectivity (degree and closeness centrality). In our experiment, participants undertook an explicit graph-learning task, and the next day task-based fMRI was acquired that allowed comparing representations in the brain with models derived from the graph. For this comparison, we employed Representational Similarity Analysis (RSA;(Kriegeskorte et al., 2008)), which allows comparing brain activation patterns to different models of the task and thus allows identifying which brain area represents which task feature.

## Methods

### Participants

Twenty-five healthy young men aged between 18 and 35 years (25.40 ± 7.03) took part in the study (the sample was filled if dropouts occurred until the final sample was reached). As for our previous work (Feld et al., 2022), we chose to only study men, since women differ in their learning across the menstrual cycle (Brown et al., 2023; Postma et al., 1999). This decision was a bid to study a homogenous sample with low noise thereby enhancing our chances of detecting a meaningful difference between our experimental conditions (Nebe et al., 2023). Participants were non-smokers, fluent in English, not currently under medication and did not have any physical or mental disorders. They all reported having a regular sleep schedule, going to bed before midnight and waking up before 8:00 am. In addition, participants did not work night shifts and were not diagnosed with sleep disorders and they did not travel across time zones. Finally, they did not report any stressful events such as exams or deadlines before or during the experiment. The experiment was approved by the UCL ethics committee (ID number: 8951/002) and all research was performed in accordance with relevant guidelines and regulations. Written informed consent was obtained from each participant before starting the experiment. Participants were compensated financially for their participation.

### Procedure

The experiment consisted of a single session that was divided into a learning phase and a retrieval phase (see Figure 1A for a timeline). During the learning phase up to three participants were invited, scheduled each one hour after the other between 8 pm and 10 pm. Here, they learned the graph task, performed the Psychomotor Vigilance Task (PVT, Dinges et al., 1997) and then answered the Stanford Sleepiness Scale (SSS, Hoddes et al., 1973) and the Positive and Negative Affect Schedule (PANAS, Watson et al., 1988). They then returned home to sleep and returned to the lab 12 hours later. Hence, between 8 am and 11 am they took part in the MRI scanning and afterwards performed the retrieval of the graph outside of the scanner. Next, PVT, SSS and PANAS were assessed again. Finally, participants filled in the Santa Barbara Sense-Of-Direction scale (SBSOD, Hegarty et al., 2002) asking questions about spatial and navigational abilities and completed the Navigational Strategies Questionnaire (NSQ, Brunec et al., 2019) asking questions about their experiences with navigation and their navigation strategy. See table 1 for descriptive data on control measures.

**Figure 1.**
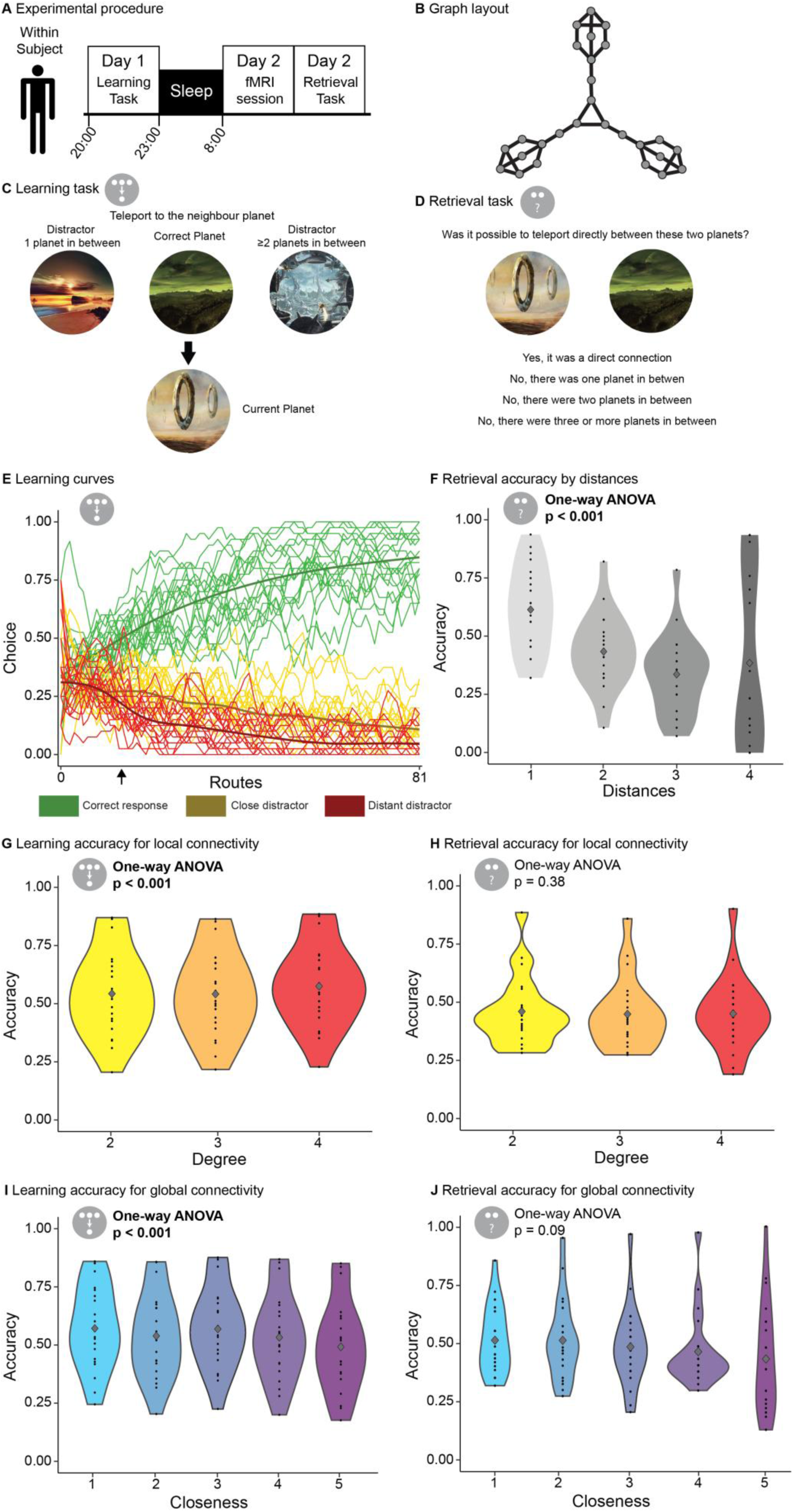
procedure and memory tasks. A) Participants took part in a single session spanning approximately 14 hours. They learned the graph task in the evening and fMRI data were collected in the morning followed by memory retrieval. B) The graph layout was selected to orthogonalize degree centrality and closeness centrality. It was never shown to the participants. C) Rather, the learning task consisted of a three alternatives forced choice paradigm, where participants learned that it was possible to teleport from one planet to another following the structure of an underlying graph. D) At retrieval, participants indicated the distance between planets on the graph. E) learning curves of all participants (thin lines) and their rolling mean (thick curves, n=5). Correct responses in green, close distractor in yellow and distant distractor in red. The black arrow indicates when participants’ performance was significantly biased by the graph structure (i.e., accuracy of the close and distant distractor started to differ according to a paired t-test). F-G) mean accuracy (large dot) and individual accuracy (small dots) per distance at learning and retrieval sessions for distances, local and global connectivity, respectively. Shaded area indicates the smoothed density.

**Table 1.** Descriptives of control measures.

|  | Learning |  | Retrieval |  |
| --- | --- | --- | --- | --- |
|  | mean | SD | mean | SD |
| PVT (reaction speed) | 3.14 | 0.27 | 3.24 | 0.26 |
| SSS | 2.71 | 1.00 | 2.57 | 0.87 |
| PANAS positive | 3.16 | 0.68 | 3.01 | 0.84 |
| PANAS negative | 1.55 | 0.50 | 1.30 | 0.29 |
| Mapping tendency | NA | NA | 2.30 | 4.78 |
| SBSS | NA | NA | 4.74 | 1.13 |

### Graph structure

We relied on the same graph structure that we previously used to show that sleep selectively enhances nodes of higher local and global importance (for details see (Feld et al., 2022), for a top down view of the network see Figure 1B). Briefly, the graph consisted of 27 nodes and 36 edges. Each node was a picture of an alien landscape and edges were connections learned in the learning task. Importantly, node identities (pictures) were randomized across participants and the graph was structured to orthogonalize degree centrality and closeness/betweenness centrality allowing us to investigate these two features and their representation in the brain independently. Note, in the methods and results we refer to closeness and degree centrality when talking about node features, but more broadly use the terms local and global connectivity in the introduction and discussion sections. To keep the same procedure as in the original study, we also rewarded some nodes in a different stage of the task, but did not analyse those data here. Note, in our previous study the reward procedure had no effect on memory for the nodes. Importantly, participants never saw the graph structure, but had to infer it from the learning trials and participants did not report using mapping strategies in the debriefing questionnaire. We programmed the graph tasks in Matlab, using the Psychophysics Toolbox extension (Brainard & Vision, 1997).

### Learning and retrieval tasks

Learning and retrieval followed were performed outside the scanner on desktop computers in the behavioural experimentation facilities of the Department for Psychology at UCL and integrated into a cover task that gamefied the experience (details can be found in Feld et al., 2022). This ensured sustained motivation for the 1.5 hours learning task. Before learning, participants were familiarized with the planets, which were shown together with a made up name on the centre of the screen (2 s each, ITI 0.5 s). During learning (Figure 1B), participants were shown a current planet as well as a choice of three other planets. They were asked to choose which planet it was possible to teleport to (three alternatives forced choice, see (Feld et al., 2022; Kern et al., 2024) previous use of this learning paradigm). One of the planets was the correct answer and two were incorrect distractors (one close distractor that was two edges away from the current planet and one distant distractor that was three or more edges away on the shortest path). After choosing, the participants received feedback, the correct planet moved to the current planet location and a new trial started. The participants travelled through the graph on eight such transitions forming one route. Routes were chosen carefully to balance exposure to transitions across the learning experience. For this, all possible 8 step routes without repetition of a transition of the graph were calculated. Then the first route was chosen randomly from this set and the next two routes were chosen to be the mirror symmetric counterparts of this route. We then proceeded to choose the next route from the set of routes that balanced the amount of exposure best between all transitions. Again, the mirror symmetric counterparts were added and the procedure was repeated until 81 routes were allocated. This led to a balancing of transitions across the learning and between participants meaning that transitions were experienced equally often over the whole sample (note, this means high degree nodes were experienced more often). Hence, the learning consisted of a total of 81 routes, i.e., participants performed 648 choices. They had 10 s for each choice. During retrieval, we presented all possible combinations of two planets (351 trials) and asked the participants to choose whether they were directly connected, two, three or more edges apart (see Figure 1C). In separate tasks, we asked about the planet names and reward contingencies (data not shown).

### fMRI task

Participants were instructed on the fMRI task in the room adjacent to the scanner and performed training trials on a laptop until they had understood the task. In the scanner, the head coil was equipped with a mirror that allowed viewing a monitor at the far end of the scanner that was mirrored from a computer running in the operator’s room. The task was gamified in line with the overall story and participants were told to look out for diamonds that they must scavenge from the planets that made up the graph. We chose only to include 15 of the 27 nodes to balance the features of interest and reduce the amount of trials (Figure 2A, black nodes). In addition to the pictures of planets from the graph, we added three completely new pictures, which were added to the GLM below to account for brain activation unrelated to the memorized graph. In order to show each possible transition between these 18 nodes, we used a de Bruijn sequence, i.e., a sequence in which every possible transition occurs exactly once as a substring (e.g., if we had 3 nodes 1,2,3,1,3,2,1 would fulfil this constraint), which led to 324 trials per run (approximately 11 minutes, 4 runs in total). Each image was shown for 1.5 s with a mean ITI of 800 ms (minimum 250 ms, maximum 12000 ms) drawn from a gamma distribution (Figure 2B). On each trial a small distortion (white noise) was presented randomly within the left or to the right part of the shown planet and participants had to detect it and respond with a button press using the index or the middle finger of the right hand for left or right responses, respectively. Each participant repeated four runs of the fMRI task.

**Figure 2:**
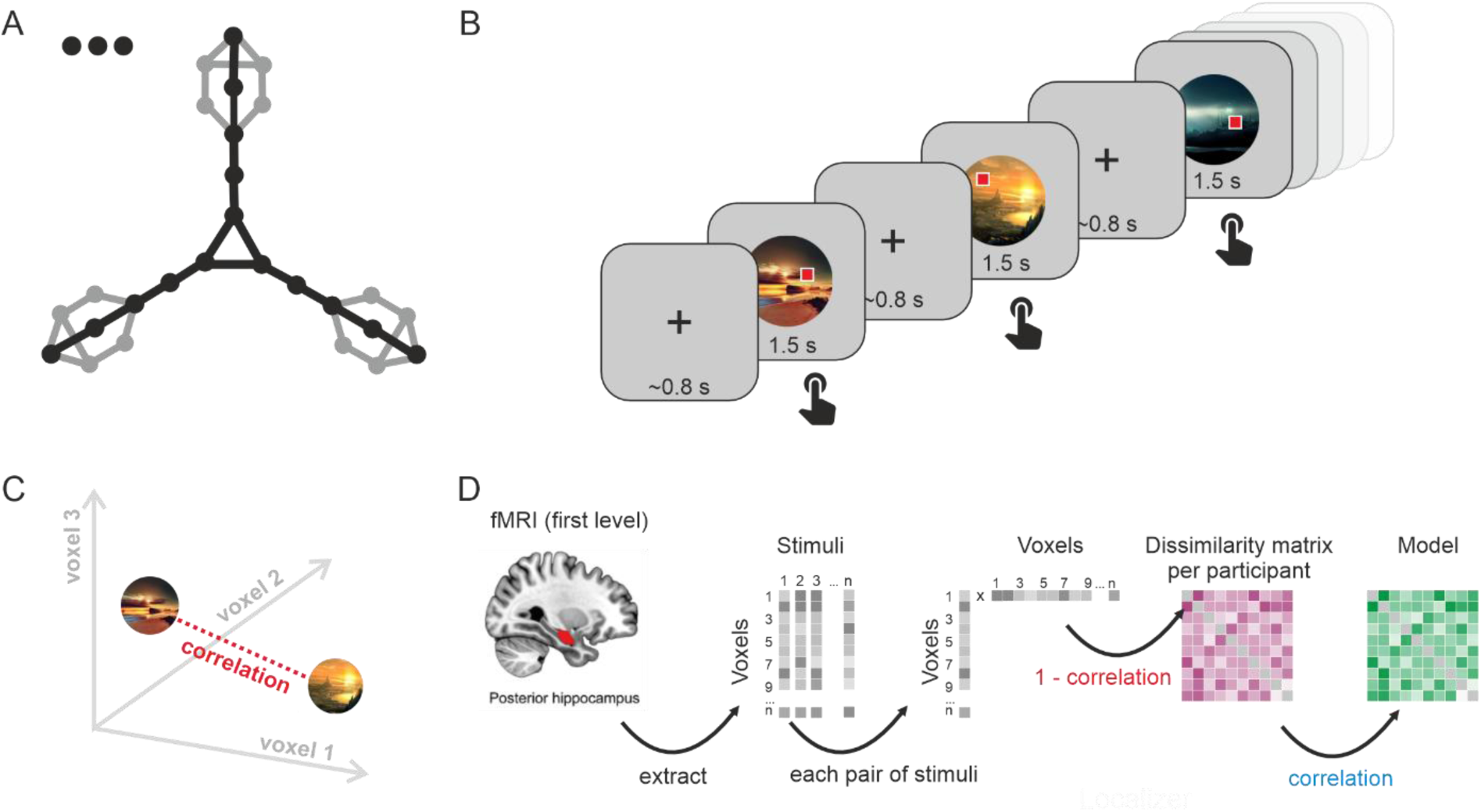
fMRI task and data processing. A) For the fMRI-task, we used a set of 15 nodes from the graph (black), i.e., nodes from the distal arms that had redundant topological features were omitted (grey), to reduce the number of stimuli shown. We added three unconnected nodes to capture variance related to stimulus presentation without a memory component. B) Each image representing a node was shown to the participant 18 times in a de Bruijn sequence, i.e., each possible transition of a completely connected graph (not the learned graph) was shown once. The task was repeated four times. C) Data analysis relied on RSA, which conceptually captures the similarity of two stimuli in a multidimensional voxel-space (three voxels shown for visualisation). D) For this, we first ran a first level fMRI analysis to receive beta coefficients for each stimulus for all voxels. We next extracted the voxels of an ROI (e.g., posterior hippocampus) and restructured them to a vector per stimulus. Each possible stimulus pair was then correlated producing a dissimilarity matrix (DM) per participant and ROI (distance = 1-r) that could then be compared to model DMs (see Figure 3 for ROIs and model DMs as well as methods section for details).

### MRI data acquisition and preprocessing

MRI scanning was conducted on an 1.5T Siemens Avanto scanner at University College London. Anatomical MPRAGE images were acquired with 1 mm isotropic resolution, TR=2.73 s, TE=3.57 ms, and flip angle 7°. Functional gradient echo EPI images were acquired in 32 slices with 3.2 mm thickness, 3.2 mm gap, 3.2x3.2 mm in-plane resolution, TR=1.42 s, TE=45 ms, multi-band factor 2, and flip angle 75°.

MRI data preprocessing was conducted with SPM 12 (v7738) running in Matlab R2020a. Anatomical images were segmented and normalized to the SPM 12 TPM MNI template. Functional images were slice-time corrected, realigned to the mean image, coregistered to the anatomical image, rescaled to 3mm isotropic resolution, normalized by applying the forward deformation field estimated from the anatomical image and smoothed by a FWHM=4x4x4 mm Gaussian kernel. Data from 4 participants was excluded from further analysis due to excessive movement (criteria: >3mm translation or >3°rotation volume-to-volume; >25% motion-affected volumes). For first level analyses, onset of node presentations were modelled with stick functions convolved with the SPM 12 canonical hemodynamic response function and entered into a multi-session general linear model that further contained the 6 standard movement regressors, white matter and cerebrospinal fluid nuisance signals and dummy regressors marking volumes affected by small movements (framewise displacement FD>0.5 mm, global frame-to-frame intensity change z>4). A high-pass filter with a cut-off of 128 s was applied. For each node, a contrast against implicit baseline over all sessions was estimated and used for further analyses.

### Representational Similarity Analysis

RSA is a multivariate analysis that works by extracting brain activity of a volume of voxels, either raw BOLD activity or beta weights from a task-based GLM (see Figure 2C and 2D). The voxel volumes from several items are each unfolded to a vector and related to each other, generating an empirical dissimilarity matrix (DM). Dissimilarity matrices from several candidate models (see Figure 3 A) are statistically related to the empirical DMs from different volumes. A brain area represents task-relevant information if the model DM and the empirical DM are related more strongly than chance would suggest. In this way, we are able to compare graph-derived models with the fMRI data to show which brain areas are tracking which connectivity properties of the abstract knowledge network under study.

**Figure 3.**
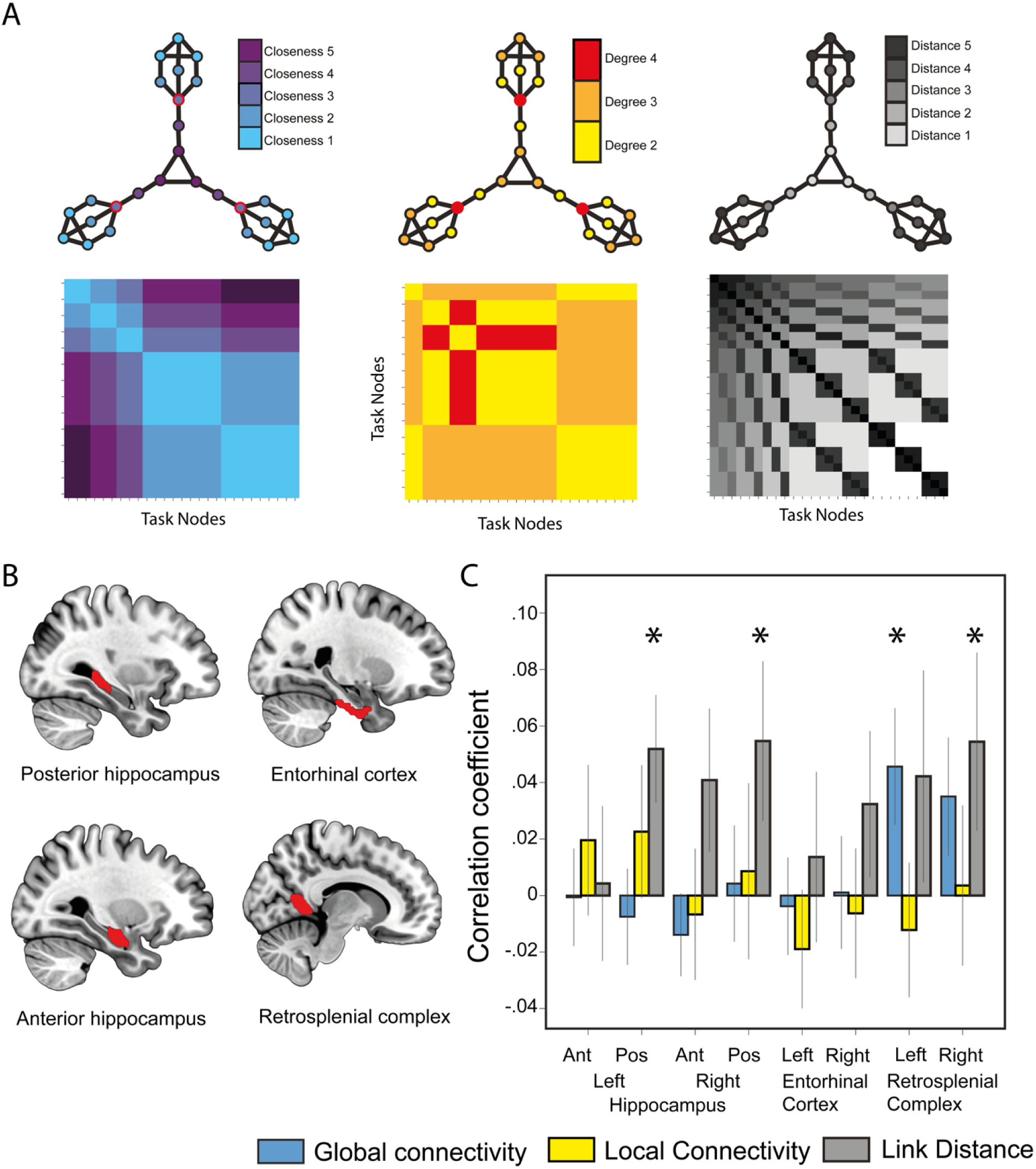
graph topologies and model DMs. A) In the top panel, the graph is shown with different topologies. On the left, degree centrality of each node, i.e., the number of edges it has. In the middle, closeness centrality, i.e., the cumulative shortest distance to any other node on the graph. Note that we chose closeness centrality here, as it was highly correlated to betweenness centrality in our graph and thus no two independent models could be used. On the right, distance from the center via the shortest path (note that each node pair has an individual distance not depicted here). In the bottom panel, the DMs of the corresponding graph topologies is shown. I.e., for degree and closeness centrality we calculated the difference between each node pair (i.e., if an edge connects a degree three node with a degree two node the DM would be filled with a one for that transition). For distance, we used the shortest path between each node. B) Brains show the ROIs used to extract voxels for the RSA analysis. C) The relationship between the model DRMs and the empirical DRM of each ROI volume. We found robust relationships between model RDMs and brain activity for distance in the lefts and right posterior hippocampus and the right RSC as well as for closeness centrality in the left RSC. Stars indicate significant relationships (p ≤ 0.05).

We defined anatomical masks for bilateral posterior hippocampus, entorhinal cortex, and retrosplenial complex (RSC) based on previous work implicating these regions in graph and spatial representation (Garvert et al., 2017; Peer & Epstein, 2021; Schapiro et al., 2013). Hippocampal masks were derived from the AAL3 atlas (Rolls et al., 2020) and divided into anterior and posterior portions at the midpoint along the anterior–posterior axis. Entorhinal cortex masks were extracted from the HCPex atlas (Huang et al., 2022). RSC masks were obtained from the Julian scene-selective parcels (Julian et al., 2012). All masks were resampled to functional space using SPM12, with voxels included if interpolated intensity exceeded 0.1. Final ROI sizes were: left/right posterior hippocampus (118/113 voxels), left/right entorhinal cortex (77/75 voxels), and left/right RSC (245/331 voxels). Statistical significance was assessed using one-sample, one-tailed t-tests (α = 0.05), testing whether mean model–brain correlations were significantly greater than zero. As each ROI×model comparison was tied to a specific a priori directional hypothesis, we report uncorrected one-tailed p-values.

### Behavioural Analysis

To obtain the learning results for each distance, we used the same methods as Feld et al. (2022). For this, we took into account the learning data by weighting each edge of the graph. For distance 1, since participants saw a transition 8 times, the weight for the first time was 1/8, second time 2/8 and last time 8/8. This weighting was applied to account for the last exposures giving a more precise estimate of current performance than the first ones. Then, we obtain a single accuracy value for each transition by multiplying the weight by the accuracy (1/0), summing it together and dividing it by the sum of all the weights. For distances 2 to 4, we used the graph structure to calculate performance between two given nodes. Specifically, the weighted accuracies of the distance 1 edges along the path between the two nodes were multiplied. We chose to multiply rather than add the accuracies, as the sum otherwise could be larger than 1 and multiplication takes into account that inferring across several edges requires knowledge about each edge in between (i.e., if the participants gets A-B right in 50% of cases and B-C right in 50% of cases, we would expect them to correctly report A-C in 25% of cases). The retrieval data were calculated with the hit rates for distance 1 to distance 4, i.e., all trials of a specific distance were averaged irrespective the specific nodes. The results for each node feature (i.e. local and global connectivity) were computed by averaging the measured data (learning or retrieval accuracies) for each feature. Hence, each node with a centrality of interest (e.g., all nodes with a value of closeness centrality of 2) and the judgements about its relation to all the other nodes of the graph were included into the average. The statistical analysis relied mainly on repeated measures ANOVA and paired t-tests.

## Results

### Behavioural results

Results from the learning session were comparable to Feld et al. (2022) which used the same task. Participants learned to distinguish the correct answer from the distractors with an average of 82.38% correct in the last third of the learning trials. As in our previous work, participants chose the close distractor significantly more frequently than the distant distractor after learning 16 routes indicating that they inferred the graph structure (Figure 1E). For the retrieval results, when separating the results by distances, participant more accurately indicated the distance between two nodes the closer the nodes were together on the graph (one-way ANOVA F(1,19)=9.78 ; p<0.001) (Figure 1F).

Regarding graph topology, we found that during the learning session participants made more correct choices for nodes that had higher degree (one-way ANOVA F(1,19)=54.56; p<0.001, Figure 1G). In contrast, participants made more errors for nodes that had higher closeness (one-way ANOVA F(1,17=34.88 ; p<0.001) (Figure 1H). In the retrieval session, there was no evidence of an effect of degree on memory performance (one-way ANOVA F(1,19)=1.01 ;p=0.38, Figure 1H). Whereas, there was a trend towards participants having worse memory performance for nodes with high closeness (one-way ANOVA F(1,17)=2.40 ;p=0.09, Figure 1J).

### Representational Similarity Analysis

Our main question concerned how graph topological information is represented by the human brain. Three candidate graph metrics were formalised as model dissimilarity matrices (RSA; Figure 3A) and subsequently compared to dissimilarity matrices of BOLD activity based on multivoxel patterns elicited by each node in the graph per anatomical region of interest. As detailed in the methods, the task environment was designed to dissociate contributions of local versus global properties of the learned network structure.

Previous work has shown functional associations with graph- and space-related distance information in distinct brain areas including the hippocampus (Schapiro et al., 2013), entorhinal cortex (Garvert et al., 2017), and RSC (Peer et al., 2021). Based on this prior work, we defined a priori anatomical masks for these regions (Figure 3B). Note that our anatomical mask of the RSC includes voxels identified by previous work to functionally behave as expected for the retrosplenial cortex, and thus extends beyond masks restricted to retrosplenial cortex proper.

Our RSA analysis (Figure 3C) identified significant representation of graph distance in bilateral posterior hippocampus (left: t(20)=2.73, p=0.006; right: t(20)=1.94, p=0.033) and in the right RSC (t(20)=1.73, p=0.049). Across our anatomical ROIs we found no significant representation of degree centrality (local connectivity). However, closeness centrality (global connectivity) was significantly represented in left RSC (t(20)=2.22, p=0.019), with a non-significant trend in right RSC (t(20)=1.69, p=0.053).

## Discussion

The brain structures that support cognitive maps of physical space have been found to support maps of abstract knowledge spaces as well (Epstein et al., 2017; Garvert et al., 2017). Here, we use a graph-based learning paradigm to study the brain regions involved in memory representations of a newly learned abstract knowledge space. By orthogonalizing the global and local centrality features of the network structure in which this abstract knowledge was organised, we were able to disentangle which brain regions were involved in representing global and local properties of the network. We found that link distance was represented in the posterior hippocampus and the right RSC. We found no representation of local connectivity (degree centrality – how many direct neighbours a node has). We found that global centrality measures (closeness/betweenness centrality) were represented in the left RSC.

Our results add to the emerging picture of the hippocampal formation and related areas mapping relations across social, temporal and conceptual dimensions (Bellmund et al., 2022; Constantinescu et al., 2016; Garvert et al., 2017; Tavares et al., 2015). Our findings extend this line of work by showing that the brain regions that differently represent the properties of a network structure in physical space are also involved in representing such properties for an abstract knowledge space.

Our data shows that the posterior hippocampus and the RSC are engaged in representing the link distance in a network of abstract knowledge. Both regions have previously been found to track the distance to the goal during spatial navigation (Patai et al., 2019). In an experiment in which participants had to navigate through segmented environments, activation in the hippocampus and RSC contained information about the distances between objects (Peer & Epstein, 2021). Another study found that RSC activation correlated with distance in a social network (Peer et al., 2021). And, a previous study exploring community structure in networks found hippocampal activation to be sensitive to distance (Schapiro et al., 2013), also in line with our findings. Interestingly, a study of navigation in a large, compartmentalised space, found that, while the anterior hippocampus encoded closer relations (within a room), the posterior hippocampus and the RSC encoded more distant relationships (within the wider building) (Kim & Maguire, 2018). Here, we find a similar brain activation pattern related to link distance when participants navigated through abstract, rather than spatial, dimensions.

Interestingly, we did not find any evidence for representation of graph topological features in the entorhinal cortex. In a study of abstract knowledge space, Garvert and colleagues (Garvert et al., 2017) found the entorhinal cortex to be engaged in tracking link distance, consistent with the presence of grid cells in that region (Hafting et al., 2005), which have been found to provide a metric for distances in physical space (Bush et al., 2015; Stemmler et al., 2015). One explanation for the difference in our results is that, unlike the study by Garvert and colleagues, our task involves explicit learning. A later study using that same data found that semantic distance was represented in the posterior hippocampus (Zheng et al., 2024), in line with other findings that hippocampal activity reflects distances in semantic spaces (Pacheco Estefan et al., 2021; Solomon et al., 2019). It might be that, because our task involved explicit learning, our link distance measure resembles the semantic distance measure in their paradigm. Note, that imaging the entorhinal cortex and the anterior hippocampus is technically demanding and contradicting results may therefore be explained by differences in signal to noise ratio.

Notably, the RSC was also involved in representing global connectivity (closeness/betweenness centrality). The RSC has been associated with the representation of different spatial features, as well as with the integration and updating of representations (Vann et al., 2009). An experiment in which rats had to travel through a maze with recurrent structure at multiple scales found that the RSC represented distance from any position on the maze to any other position (Alexander & Nitz, 2017). Although they were not using graph theoretical metrics in their analysis, their finding is effectively that the RSC represented global connectivity. Here, we extend this result to the representation of abstract knowledge space. Alexander & Nitz also found that RSC neurons exhibited recurrent firing patterns that defined route sub-spaces at multiple spatial scales, which implies that the RSC is involved in representing multiple spatial relationships. The RSC has reciprocal connections to several brain regions that map space using different reference frames, and the dominant view is that it enables coordinate transformation for switching between allocentric and egocentric frames of reference in navigation (Alexander & Nitz, 2017; Byrne et al., 2007). However, coordinate transformation by itself cannot explain our results. Rather, since our task involves learning a graph network from a series of paired associates, our findings indicate that the RSC may enable building a larger map from these distinct episodes by keeping track of the position within a larger network of memory contents.

Since RSC is positioned between hippocampus and neocortex, it has been suggested to play an important role for consolidation (de Almeida-Filho et al., 2021; Maviel et al., 2004; Milczarek & Vann, 2020). In physical navigation, the RSC is more active in well-consolidated environments: Patai and colleagues (Patai et al., 2019) tested participants navigating a familiar and a new environment, and they found that the posterior hippocampus tracked distance to goal in the new environment and the RSC tracked distance to goal in the familiar environment. We have previously showed that closeness centrality and degree centrality are relevance signals for sleep consolidation (Feld et al., 2022), and a separate study found that optogenetically enhancing RSC activity leads to faster consolidation during subsequent sleep (de Sousa et al., 2019). Considering our results in conjunction with previous findings, it appears that the RSC might provide a relevance signal regarding global connectivity, which serves to select which features of the environment should be prioritised during consolidation. And the RSC may play this role in consolidation both for physical and for abstract knowledge spaces.

Previous research suggests that the degree centrality of a node is represented in the posterior hippocampus (Javadi et al., 2017), in line with the view that this region tracks local, fine-grained features, while the anterior hippocampus and the RSC track global, coarse-grained features of the environment (Farzanfar et al., 2023). Also, shorter paths have been shown to be represented in the anterior hippocampus, where local connectivity such as degree centrality may be expected (Kim & Maguire, 2018). However, using RSA, we did not find any correlations with degree centrality. In the previous study that correlated degree centrality with posterior hippocampal activity, participants were actively planning their way through the network, and this may have been key to finding the previous association. Further work will be required to explore how brain regions represent centrality measures in different contexts to determine if this was a key factor.

While we managed to orthogonalize degree centrality and closeness centrality, we were not able to orthogonalize betweenness and closeness centrality. Both of these global metrics are analogous in the abstract knowledge network in our task. Future work could design a network in such a way that the two metrics came apart. This would reveal whether separate regions are tasked with representing each of the two global metrics. We did not attempt this here because such a network requires many more nodes to learn and would have involved very extended training. However, further studies with different approaches to learning may make that possible.

We set up our task so that it was not explicitly spatial, with participants only exposed to a small set of items on each trial and no mappings of the network displayed. It is possible that some participants created a map in their minds eye to solve the task. However, none of the participants explicitly reported doing this when debriefed. It would be interesting to compare our current findings to a task where participants explicitly learn the shape of the network and structure, e.g., using a network map, as would it be interesting to manipulate sleep to examine the impact of consolidation on network representations (Feld et al., 2022).

In sum, it appears that the brain structures involved in the internal mapping of networks in physical space also represent networks of abstract information. Here, by studying how the brain activity of certain regions of the brain are correlated with network properties we can understand the different contributions of these brain regions to the acquisition of integrated knowledge. Specifically, the posterior hippocampus and the right RSC represent distance, and that the left RSC represents global connectivity. Our results are consistent with the idea that the RSC keeps track of position within the extended network, enabling the building of a more global map, as well as being involved in long-term memory consolidation of such integrated information. Taken together, our research underlines the relevance of hippocampus and RSC to jointly represent a spaces even within a nonspatial, abstract domain.

## Acknowledgements

We thank Timothy E.J. Behrens and Bernhard Staresina for their valuable support of the project design.

## Funding

This research was supported by DFG grants (project numbers 404679789 and 290208383) and an ERC Consolidator Grant (MemoryTracker, grant agreement number 101170886) to GBF, a James F McDonnell Scholar award to HJS and funding by British Academy [PFSS23\230053] and by the Leverhulme Trust [LIP-2023-002] to PFV.

## Data availability

Data are available upon request.

## Notes

### Competing Interest Statement

The authors have declared no competing interest.

